## Supplemental Tables for "Developmental patterning function of GNOM ARF-GEF mediated from the plasma membrane"

Supplementary File 1. Additional materials and methods.

Supplementary File 1a. Lines generated as part of this study.

**Line name**

*gn^s^ CLC2_pro_:CLC2-GFP*

*gn^s^ LAT52_pro_:TPLATE-GFP RPS5A_pro_:AP2A1-TagRFP*

*gn^s^ secRFP*

*big3 GNOM-GFP VAN3_pro_:VAN3-mRFP*

*XVE>>amiCHCa GNOM-GFP VAN3_pro_:VAN3-mRFP*

*GNL1_pro_:GNL1^sens^-YFP gnl1* x *Col-0*

*gn^s^ GN_pro_:GN-GFP*

*gn^s^ GN_pro_:GGL-GFP*

*gn^s^ GN_pro_:LGG-GFP*

*gn^s^ GN_pro_:GLG-GFP*

*gn^s^* +/- *GN_pro_:LGL-GFP*

*gn^s^* +/- *GN_pro_:GLL-GFP*

*gn^s^* +/- *GN_pro_:GNL1-GFP*

*GN_pro_:GN-GFP*

*GN_pro_:GGL-GFP*

*GN_pro_:LGG-GFP*

*GN_pro_:GLG-GFP*

*GN_pro_:LGL-GFP*

*GN_pro_:GLL-GFP*

*GN_pro_:GNL1-GFP*

*gn^s^ GN_pro_:GN^fwr^-GFP*

*GN_pro_:GN^fwr^-GFP*

*gnl1-2 VHAa1_pro_:VHAa1-GFP*

Supplementary File 1b. Primers used in this study.

| **Primer name** | **Sequence** | **Purpose** |
| --- | --- | --- |
| gn-SALK-F | gggtatatgctccggctgta | *gn^s^* genotyping |
| gn-SALK-R | tcagaacatcctacaggaccaa |  |
| LBb1.3 | ATTTTGCCGATTTCGGAAC |  |
| big3-1-F | GCCTCCTGATGCCACTTTCT | big3-1 genotyping |
| big3-1-R | ATCATGTGCTGCTGTCGTGA |  |
| attB1-GN-F | GGGGACAAGTTTGTACAAAAAAGCAGGCTTTATGGGTCGCCTAAAGTTGC | GN, FWR, chimeras cloning |
| attB2-GN-Rns | GGGGACCACTTTGTACAAGAAAGCTGGGTCCGAACCAGTTGTGTTTTCAGG |  |
| attB1-GNL1-F | GGGGACAAGTTTGTACAAAAAAGCAGGCTTTATGGGGTATCAGAATCATCCTT | GNL1, chimeras cloning |
| attB2-GNL1-Rns | GGGGACCACTTTGTACAAGAAAGCTGGGTCGACCTCATTTCCCGGTACCG |  |
| GLG1-R | CAAAGGGAACCCAGTGATTTGGATCACTAT | chimeras cloning |
| GLG2-F | AAATCACTGGGTTCCCTTTGTCCGAAAGGT |  |
| GLG2-R | GAGTCATTTCTTGGAAGCCGGTTCCTTTAT |  |
| GLG3-F | CGGCTTCCAAGAAATGACTCCCAGCCGTTG |  |
| LGG1-R | CAAATGAAACCCAAAAATTAGGATCTCCGT |  |
| LGG2-F | TAATTTTTGGGTTTCATTTGTCCGCAGGAG |  |
| GGL1-R | CTGTCATCAGAGGAAAGCCTGCACCTTGTT |  |
| GGL2-F | AGGCTTTCCTCTGATGACAGCCAGTCGTTG |  |
| attB4-GNpro-F | GGGGACAACTTTGTATAGAAAAGTTGtctagaggtgtgtatgataatgaatattga | GN promoter cloning |
| attB1r-GNpro-R | GGGGACTGCTTTTTTGTACAAACTTGttaatctgctcaaatcttcagcc |  |
| GNfwr-F | ACAAGCTGAATTTTTGTTGCAGCTTGCGCGTGC | FWR mutagenesis |
| GNfwr-R | GCTGCAACAAAAATTCAGCTTGTAGAAACTTGC |  |

Supplementary File 1c. Constructs generated in this study.

| **Plasmid name** | **Notes** |
| --- | --- |
| GNOM /pDONR221 | without STOP codon |
| FEWERROOTS/pDONR221 | without STOP codon |
| GNOM-LIKE1/pDONR221 | without STOP codon |
| GGL/pDONR221 | without STOP codon |
| LGG/pDONR221 | without STOP codon |
| GLG/pDONR221 | without STOP codon |
| LGL/pDONR221 | without STOP codon |
| GLL/pDONR221 | without STOP codon |
| ProGNOM/pDONRP4P1r |  |
| pGN:GN-GFP/pH7m34GW |  |
| pGN:GNL1-GFP/pH7m34GW |  |
| pGN:GLL-GFP/pH7m34GW |  |
| pGN:GGL-GFP/pH7m34GW |  |
| pGN:GLG-GFP/pH7m34GW |  |
| pGN:LGL-GFP/pH7m34GW |  |
| pGN:LGG-GFP/pH7m34GW |  |
| pGN:GN^fwr^-GFP/pH7m34GW |  |
